# The Aurora B gradient sustains kinetochore stability in anaphase

**DOI:** 10.1101/2021.02.26.433106

**Authors:** Diana Papini, Mark Levasseur, Jonathan M.G. Higgins

**Affiliations:** Biosciences Institute, Faculty of Medical Sciences, Newcastle University, Newcastle upon Tyne, NE2 4HH, UK

## Abstract

Kinetochores assemble on chromosomes in mitosis to allow microtubules to attach and bring about accurate chromosome segregation. The kinases Cyclin B-Cdk1 and Aurora B are crucial for the formation of stable kinetochores. However, the activity of these two kinases appears to decline dramatically at centromeres during anaphase onset, precisely when microtubule attachments are required to move chromosomes towards opposite poles of the dividing cell. We find that, although Aurora B leaves centromeres at anaphase, a gradient of Aurora B activity centred on the central spindle is still able to phosphorylate kinetochore substrates such as Dsn1 to modulate kinetochore stability in anaphase and to regulate kinetochore disassembly as cells enter telophase. We provide a model to explain how Aurora B co-operates with Cyclin B-Cdk1 to maintain kinetochore function in anaphase.

## Introduction

The mitotic spindle acts to segregate chromosomes accurately into daughters during cell division. To accomplish this, kinetochores are built on centromeres in early mitosis to enable chromosomes to capture microtubules. Accordingly, the mechanisms that ensure that kinetochores assemble and attach to microtubules emanating from a single pole have been extensively studied. For example, in early mitosis, the kinase Aurora B plays a vital role in both kinetochore assembly and in destabilizing incorrect kinetochore-microtubule attachments^1-7^, and Cyclin B-Cdk1 is also required for key proteins to localize to the kinetochore^8-12^. In many respects, these activities serve as a prelude to the most obvious function of kinetochores: driving chromosome movement to opposing spindle poles in anaphase, when kinetochore-spindle attachments are known to be stable^13-15^. However, it is striking that key processes that control kinetochore assembly and function are driven by the activity of Cyclin B-Cdk1, which strongly declines at the metaphase to anaphase transition^16,17^, and Aurora B, which dissociates from centromeres and transfers to the central spindle in anaphase^18^. For example, the phosphorylation of the KMN (Knl1 complex, Mis12 complex, Ndc80 complex) protein Dsn1 at S100 and/or S109 by Aurora B is required for the Mis12 complex (Mis12C) to stably associate with the constitutive centromere-associated network (CCAN) proteins CENP-C and CENP-T^3,5-7,12,19-21^. However, it is widely believed that the function of Aurora B at kinetochores ceases in anaphase^15,22-24^. This raises the question of how kinetochore stability is maintained in anaphase, precisely when spindle attachments are critical to move chromosomes. A related and also understudied question is how kinetochores are disassembled as cells enter telophase.

Here we find that the previously reported gradient of Aurora B activity centered on the spindle midzone in anaphase^25^ is able to phosphorylate kinetochore substrates to regulate kinetochores in late mitosis.

## Results

### The KMN protein Dsn1 is phosphorylated at kinetochores in early anaphase

In anaphase, Aurora B focused at the central spindle generates a gradient of activity that results in high phosphorylation of substrates at the midzone and progressively lower phosphorylation towards the poles^25^. As previously reported, this gradient can be observed in HeLa cells when visualizing phosphorylation of Histone H3 at S10 (H3S10ph), a product of Aurora B activity on anaphase chromosomes, particularly when cells were treated with MPS1-IN-1, an inhibitor of the mitotic checkpoint kinase Mps1^26^, to induce lagging chromosomes (Figure S1a). A gradient of phosphorylation of the centromeric Aurora B target CENP-A S7 was also apparent in anaphase in RPE1 cells (Figure S1b), consistent with the previously observed gradient of phosphorylation on an artificial substrate targeted to centromeres with CENP-B^25^. However, despite previous speculation^2^, whether there is a gradient of Aurora B activity that influences kinetochores themselves, and whether this has functional consequences, has not been explored.

When examining the Aurora B target residue S109 on the kinetochore protein Dsn1 by immunofluorescence microscopy in HeLa cells, we found that phosphorylation was observed on essentially all prometaphase kinetochores and that this declined slightly, but clearly remained present, during metaphase (Figure 1a,b) as previously reported^4,27,28^. This contrasts with phosphorylation of a number of Aurora B target sites on Knl1 and Hec1/Ndc80 that decrease substantially during metaphase^4,28-30^. In addition, we found that Dsn1 S109 phosphorylation (Dsn1 S109ph) persisted into anaphase (Figure 1a,b). When HeLa cells were treated with MPS1-IN-1 to induce lagging chromosomes, a clear gradient of Dsn1 S109ph was revealed: kinetochores near the cell equator could be more strongly phosphorylated than those near the poles (Figure 1c,d). Linear regression confirmed that the observed gradient was significantly different from zero (p < 0.0001, F test). A similar gradient could be observed in RPE1 cells (Figure S1c,d). Dsn1 S109ph gradients were also observed with a second Dsn1ph S109 antibody (Figure S1a), and in the absence of MPS1-IN-1 in HeLa cells (see below) and in RPE1 cells (Figure S1e).

**Figure 1:**
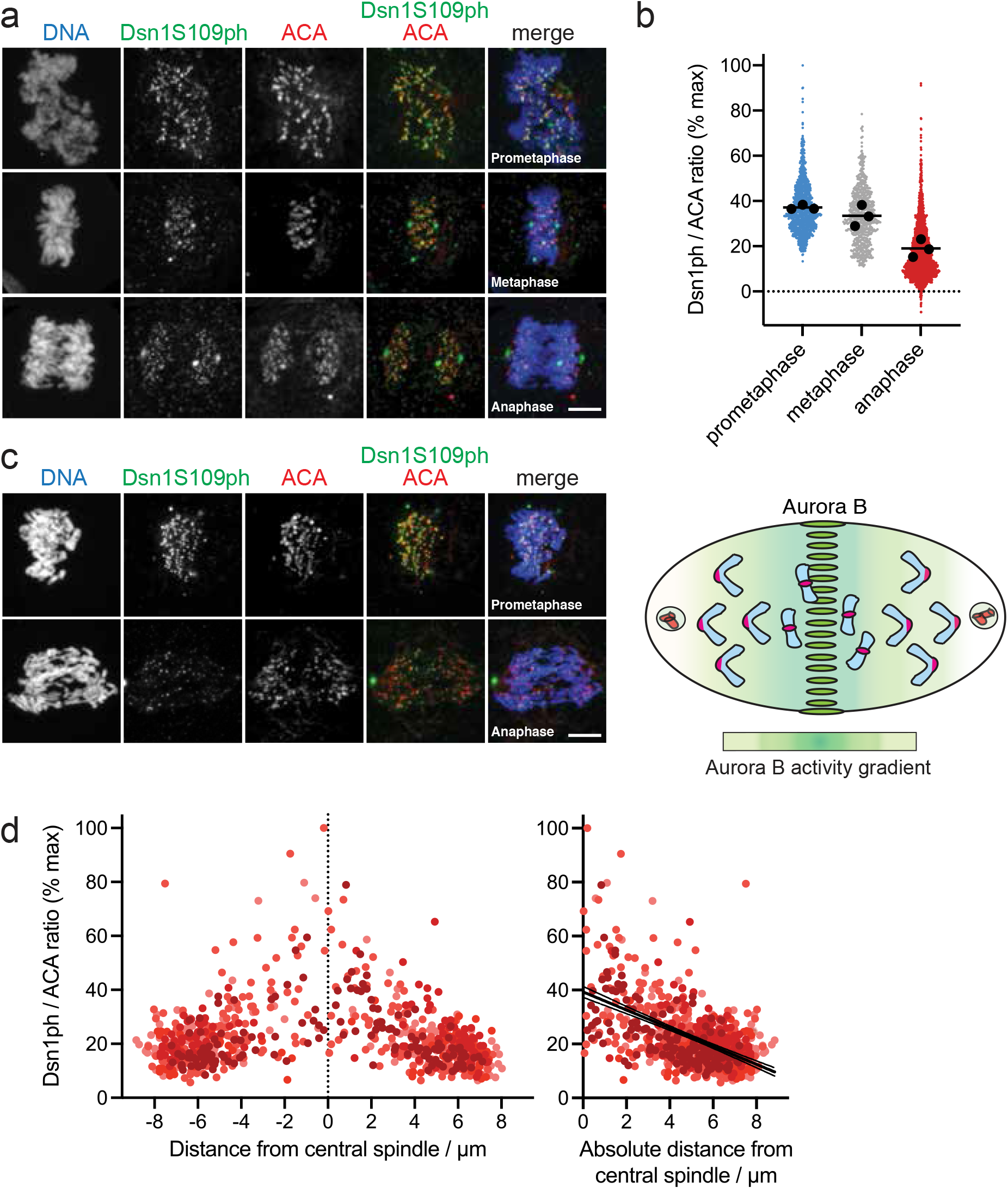
A kinetochore Aurora B substrate shows a gradient of phosphorylation in anaphase. **a**. HeLa cells were stained for DNA (blue), Dsn1S109ph (green) and ACA (red). **b**. Quantification of Dsn1S109ph in prometaphase (13 cells), metaphase (9 cells) and anaphase (23 cells) from 3 independent experiments performed as in (a). **c**. HeLa cells treated with 10 μM MPS1-IN-1 to generate lagging chromosomes that report the gradient activity of Aurora B (see diagram) were stained as in (a). Scale bars, 5 μm. **d**. Quantification of Dsn1S109ph at kinetochores as a function of distance from the central spindle in 10 cells treated as in (c), shows a gradient of phosphorylation. Using linear regression, slope = −3.4 and is non-zero (p < 0.0001; F test). Confidence intervals (95%) shown as fine lines.

### Anaphase phosphorylation of Dsn1 requires the central spindle-dependent gradient of Aurora B activity

Because Aurora B dissociates from chromosomes in anaphase, we sought to determine if the observed gradient of kinetochore Dsn1 phosphorylation was due to Aurora B activity emanating from the central spindle. First, we studied the effect of acutely interfering with Aurora B location and activity in anaphase. HeLa cells were released from a nocodazole-induced mitotic arrest into medium containing MPS1-IN-1 to produce lagging chromosomes. When nocodazole was re-added to anaphase cells for 5 min to acutely depolymerize microtubules and prevent Aurora B localization to the central spindle (Figure S2a), the gradients of both H3S10ph and Dsn1 S109ph were lost (Figure 2a). Similarly, when Aurora B kinase activity was inhibited by adding the Aurora B inhibitor ZM447439 for 5 min (Figure S2b), the gradients of both H3S10ph and Dsn1 S109ph on anaphase chromosomes were again eliminated (Figure 2b). This finding was confirmed when Dsn1 S109ph was quantified in similar experiments in which ZM447439 was added for 10 min (Figure 2c) or 3 min, both with or without MPS1-IN-1 treatment (Figure S2c,d).

**Figure 2:**
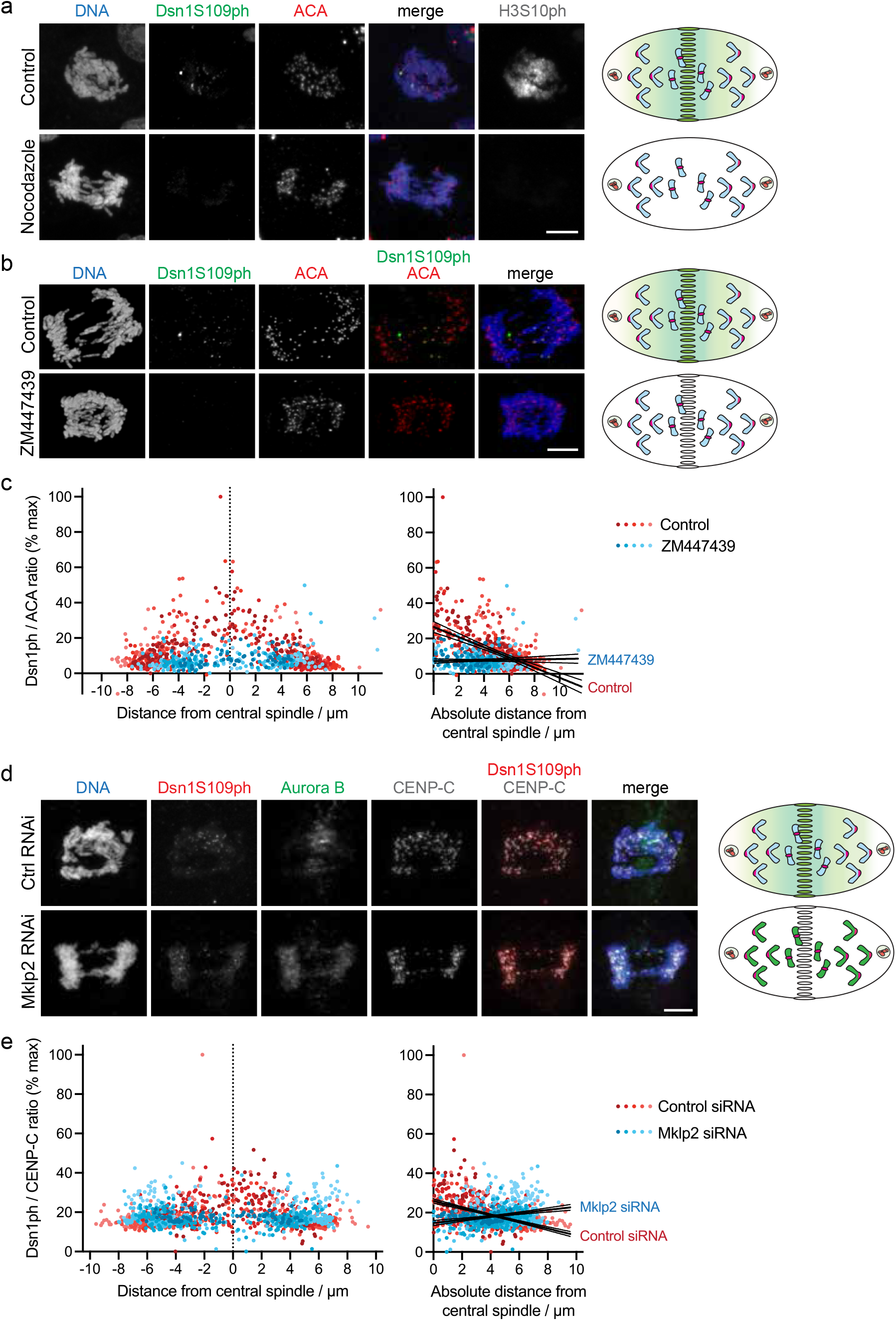
The gradient of Dsn1S109ph is dependent on central spindle Aurora B. **a**. Central spindle depolymerisation (see diagram) eliminates the Dsn1S109ph gradient. HeLa cells were enriched in anaphases with lagging chromosomes (see Methods), and nocodazole was added 5 min prior to fixation and staining for DNA (blue), Dsn1S109ph (green), ACA (red) and H3S10ph (gray). **b**. Acute Aurora B inhibition eliminates the Dsn1S109ph gradient. HeLa cells were treated with **b**. MPS1-IN-1, and then with 10 μM ZM447439 for 15 min, prior to fixation and staining for DNA (blue), Dsn1S109ph (green), and ACA (red). **c**. Quantification of Dsn1S109ph at kinetochores as a function of distance from the central spindle in 9 control and 5 ZM447439-treated cells treated as in (b). Using linear regression, for control cells, slope = −2.9 and is non-zero (p < 0.0001; F test). For ZM447439-treated cells, slope = 0.12 and is not significantly different from zero (p = 0.48). The slopes are significantly different from one another (p < 0.0001, F test). **d**. Mklp2 RNAi prevents transfer of Aurora B to the central spindle (see diagram) and weakens the Dsn1S109ph gradient. Control and Mklp2-depleted cells were treated with MPS1-IN-1 and stained for DNA (blue), Dsn1S109ph (red), Aurora B (green) and CENP-C (gray). Scale bars, 5 μm. **e**. Quantification of Dsn1S109ph at kinetochores as a function of distance from the central spindle in 8 control and 7 Mklp2-depleted cells as in (d). Using linear regression, for control cells, slope = −1.7 and is non-zero (p < 0.0001; F test). For Mklp2-depleted cells, slope = 0.83 and also non-zero (p < 0.0001; F test). The slopes are significantly different from one another (p < 0.0001, F test). Confidence intervals (95%) shown as fine lines.

Next, we depleted Mklp2 using RNAi, which prevents the normal transfer of Aurora B to the spindle midzone and leads to its retention on chromosomes^31^, and examined the gradient of Dsn1 S109ph in MPS1-IN-1-treated HeLa cells. As expected, Aurora B was largely retained on chromatin and, in addition, kinetochore Dsn1 S109 no longer showed a clear gradient of phosphorylation. Instead, a relatively equal low level of phosphorylation at kinetochores across the cell was observed (Figure 2d,e). Similar results were obtained in the absence of MPS1-IN-1 (Figure S2e), and in RPE1 cells (Figure S2f). In a final approach, we depleted PRC1 from HeLa cells to remove central spindle Aurora B, leaving it predominantly at the equatorial cortex^32^. In this situation, only kinetochores near Aurora B at the cell cortex were phosphorylated at Dsn1 S109 (or CENP-A S7; Figure S2g,h). Together, these results suggest that Dsn1 S109 phosphorylation is dependent on the gradient of Aurora B activity centered on the spindle midzone in anaphase.

### Dsn1 is found in a gradient at anaphase kinetochores

Two mechanisms each maintain up to 50% of Mis12C (containing Dsn1) at kinetochores prior to anaphase: binding to CENP-C or CENP-T^11,33^. The phosphorylation of Dsn1 at S100 and/or S109 by Aurora B is required for the stable association of Mis12C with both CENP-C and CENP-T^3,5-7,21^. Dsn1 S100/S109 phosphorylation relieves the inhibitory effect of a basic region of Dsn1 on the interaction with CENP-C and CENP-T^7,12,19-21^. The interaction of Mis12C with CENP-T, but not with CENP-C, is additionally dependent on phosphorylation of CENP-T by Cyclin B-Cdk1^8-12,21^. To determine whether, like Dsn1 S109ph, total Dsn1 is also found in a gradient, we compared the distribution of phosphorylated and total Dsn1 in HeLa cells undergoing normal anaphase. As expected^3,6,7,9^, Dsn1 persisted at kinetochores in anaphase, though it declined as anaphase progressed (Figure 3a,b). In anaphases of MPS1-IN-1-treated cells, a gradient of Dsn1 localization intensity could also be observed, including in single cells, suggesting that Dsn1 localization to kinetochores is spatially regulated (Figure S3a,b). Interestingly, however, the gradient of total Dsn1 at anaphase kinetochores was shallower than that observed for Dsn1 S109ph, meaning that Dsn1 appeared to persist at poleward kinetochores for longer than Dsn1 S109ph (Figure 3c). This suggests that there is a temporal delay in the release of Dsn1 from kinetochores following its dephosphorylation, and/or that there is a population of Dsn1 that can localize to kinetochores until late anaphase independently of Dsn1 S109 phosphorylation.

**Figure 3:**
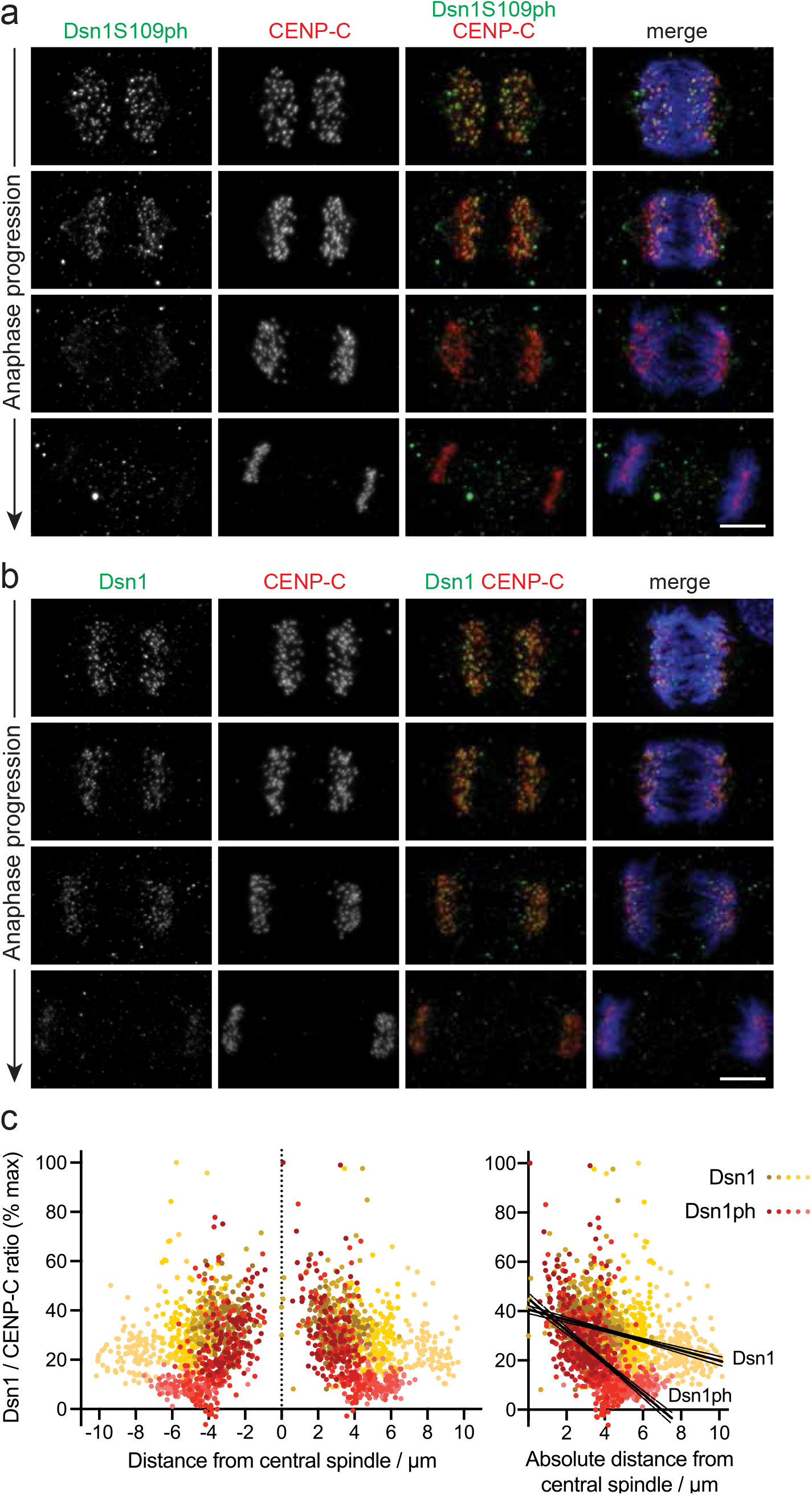
Dsn1 itself is found in a gradient at anaphase kinetochores. **a**. HeLa cells enriched in anaphases by thymidine-release were stained for DNA (blue), Dsn1S109ph (green), and CENP-C (red). **b**. HeLa cells as in (a) stained for DNA (blue), total Dsn1 (green), and CENP-C (red). Scale bar, 5 µm. **c**. Quantification of Dsn1S109ph (11 cells) and total Dsn1 (15 cells) at kinetochores as a function of distance from the central spindle. Using linear regression, for Dsn1S109ph, slope = −6.5 and is non-zero (p < 0.0001; F test). For total Dsn1, slope = −2.1 and is non-zero (p < 0.0001; F test). The slopes are significantly different from one another (p < 0.0001, F test). Confidence intervals (95%) shown as fine lines.

### Phosphorylation of Dsn1 modulates Dsn1 localization at kinetochores in anaphase

If the localization of Dsn1 to kinetochores in anaphase depends in part on its phosphorylation, as in early mitosis, then the normal distribution of Dsn1 in anaphase should require Aurora B activity. Short term inhibition of Aurora B (5 to 10 min), in either the presence or absence of MPS1-IN-1, had a minor effect on total Dsn1 signal at anaphase kinetochores in immunofluorescence experiments. The elevations/intercepts, but not the slopes, of the gradients changed with statistical significance (Figure S3c,d), suggesting a subtle effect of Aurora B inhibition on Dsn1. Consistent with this, short-term inhibition of Aurora B was also insufficient to substantially remove Dsn1 from kinetochores in prometaphase, although Dsn1 S109ph and H3S10ph were reduced (Figure S3e).

As an alternative and more tractable system to examine Dsn1 localization, we turned to fluorescent Dsn1 fusion proteins. Specifically, we compared the behavior of Dsn1-WT-GFP and a mutant in which S100 and S109 were replaced with phospho-mimetic glutamate residues (Dsn1-EE-GFP)^7^. As seen for endogenous Dsn1 (Figure 3a,b), Dsn1-WT-GFP was detected on kinetochores during anaphase, but was reduced in telophase (Figure 4a). Dsn1-EE-GFP was also present on kinetochores during anaphase (Figure 4b), but it persisted into telophase more clearly than Dsn1-WT-GFP, as previously reported^7^. Because Dsn1-GFP expression levels varied from cell to cell, it was not straightforward to use fluorescence microscopy of fixed cells to compare its localization at different stages of anaphase. Therefore, we used live cell imaging to follow the localization of Dsn1-GFP at kinetochores during anaphase in individual cells (Figure 4c). This confirmed that Dsn1-WT-GFP localization at kinetochores declined as chromosomes moved poleward in anaphase (Movie 1), as observed for endogenous Dsn1. Dsn1-EE-GFP showed a similar decline at kinetochores through anaphase (Movie 2), but was retained in telophase to a greater extent than Dsn1-WT-GFP. Short-term treatment with Aurora B inhibitor modestly, but significantly, increased the rate of Dsn1-WT-GFP dissociation from kinetochores in anaphase (Movie 3), while Dsn1-EE-GFP was less affected (Movie 4). In addition, the persistent localization of Dsn1-EE-GFP on telophase chromosomes was not greatly affected by Aurora B inhibitor (Figure 4c).

**Figure 4:**
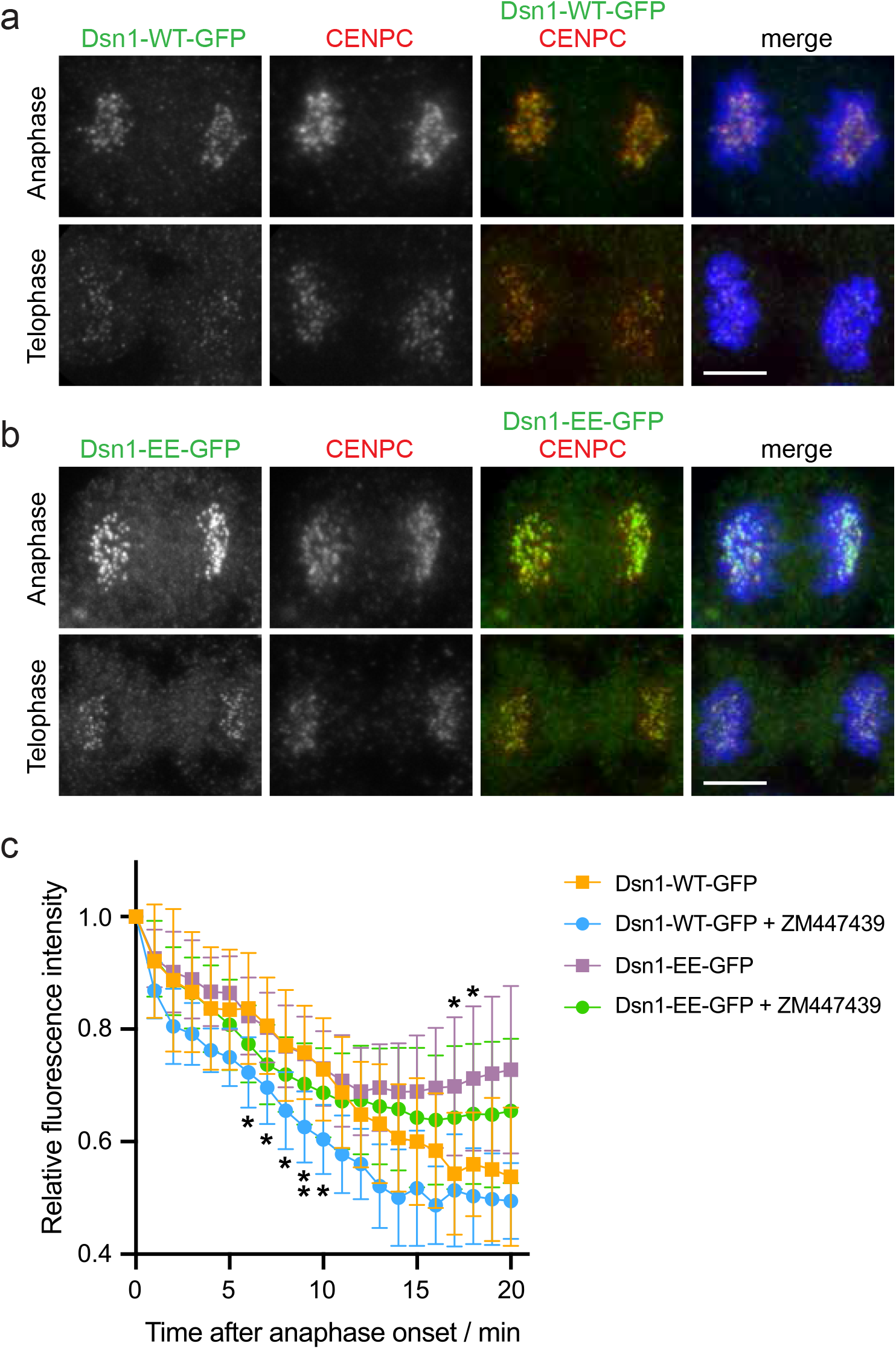
Aurora B phosphorylation modulates Dsn1 localisation at anaphase kinetochores. **a**. HeLa cells stably expressing Dsn1-WT-GFP were stained to visualize DNA (blue), GFP (green) and CENP-C (red). **b**. HeLa cells stably expressing Dsn1-EE-GFP were stained as in (a). Scale bar, 5 μm. **c**. Quantification of the kinetochore fluorescence of Dsn1-WT-GFP and Dsn1-EE-GFP in living HeLa cells imaged at 1 min intervals. Values were normalized to 1 at anaphase onset. Where indicated, cells were treated between 0 and 10 minutes prior to anaphase onset with 4 or 10 µM ZM447439. For Dsn1-WT-GFP +/-ZM447439, n = 8; and for Dsn1-EE-GFP +/-ZM447439, n = 12 and 10. * p < 0.05, ** p = 0.0085, by two-way ANOVA followed by Dunnett’s multiple comparisons test.

To explore these results further, we generated a simple mathematical model for the dissociation of Mis12C from CENP-C and CENP-T at anaphase kinetochores. We made the simplifying assumption that the dissociation can be described as a two-phase exponential decay. We made the following additional assumptions. First, at the start of anaphase, 50% of Mis12C is associated with CENP-T, regulated by both Aurora B and Cyclin B-Cdk1, and 50% is associated with CENP-C in an Aurora B-dependent manner^11,33^. Second, the dissociation of Cdk1-dependent Mis12C is likely to be faster than that of Aurora B-dependent Mis12C. Indeed, Cyclin B1 destruction reaches completion in early anaphase^17,34^, and many Cdk1-generated phosphosites are removed rapidly in anaphase^35^, including in the N-terminal region of CENP-T^8^. Third, the affinity of Aurora B-phosphorylated Mis12C for CENP-C and Cdk1-phosphorylated CENP-T is similar, as reported^21^. The model can recapitulate key features of the results, such as the similar kinetics of Dsn1-WT and Dsn1-EE dissociation from kinetochores in early anaphase (likely largely due to loss of Cdk1-dependent CENP-T binding), and the failure of approximately 50% of Dsn1 to dissociate from kinetochores in telophase when the half-life of Aurora B-dependent Mis12C is increased (Figure S4a), as seen for the Dsn1-EE mutant (Figure 4c). Fitting the model to the data in Figure 4c using non-linear regression (Figure S4b) suggested an approximate half-life for Cdk1-dependent Mis12C in anaphase of 5 min. The half-life of Aurora B-dependent Mis12C was approximately 17 min, declining to 5 min when Aurora B was inhibited (p=0.011, extra sum-of-squares F test).

These results suggested that phosphorylation of S100 and/or S109 by Aurora B is required for the normal maintenance of Dsn1 localization at kinetochores in anaphase, and that dephosphorylation is required for the complete release of Dsn1 from kinetochores in telophase.

### Dsn1 phosphorylation by Aurora B modulates kinetochore disassembly in late anaphase

In current models, the recruitment of other KMN components to kinetochores in early mitosis is partly dependent on the binding of Mis12C to CCAN proteins^36^. To determine whether Dsn1 phosphorylation modulates the localization of additional kinetochore components during mitotic exit, we examined Nuf2, a component of the Ndc80 complex (Ndc80C). Like Dsn1 itself, Nuf2 was lost from kinetochores in Dsn1-WT-GFP-expressing HeLa cells as anaphase progressed (Figure 5a,c), and kinetochore localization was low in telophase cells (Figure 5d). However, in cells expressing Dsn1-EE-GFP, the decline in kinetochore Nuf2 was less clear (Figure 5b,c) and it remained detectable at kinetochores into telophase (p < 0.0001, one-way Anova followed by Bonferroni’s multiple comparison test; Figure 5d). Inhibition of Aurora B in cells expressing Dsn1-WT-GFP compromised the Nuf2 gradient in anaphase (Figure 5c), although we noted that Aurora B inhibition also had some effect in cells expressing Dsn1-EE-GFP (Figure 5c), suggesting an additional Aurora B contribution independent of Dsn1 S100/S109 phosphorylation. The localization of Nuf2 to telophase kinetochores in Dsn1-EE-GFP expressing cells was largely resistant to Aurora B inhibitor treatment (Figure 5d), suggesting that dephosphorylation of Dsn1 S100/S109 is involved in the disassembly of the kinetochore as cells enter telophase.

**Figure 5:**
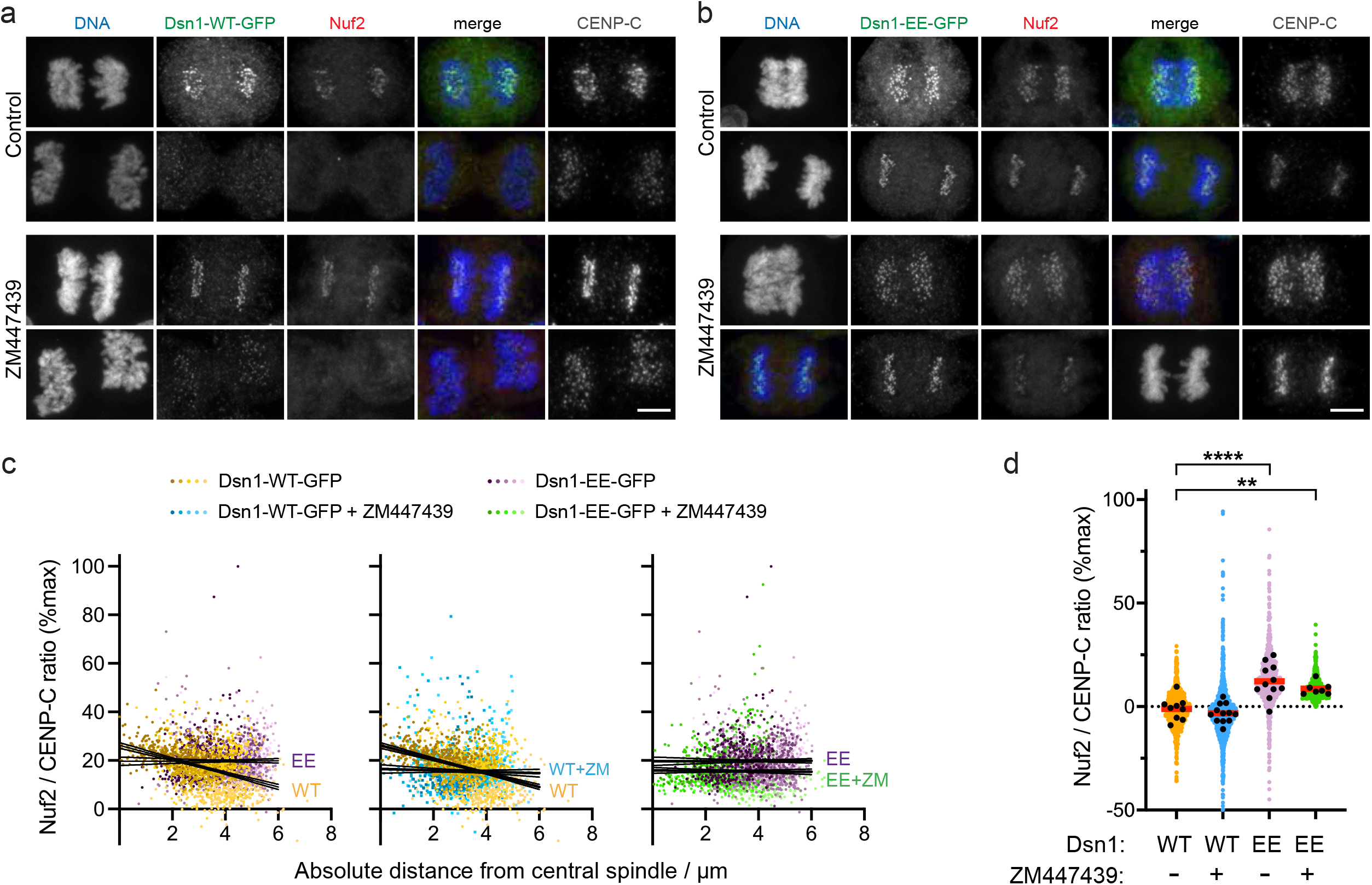
Dsn1 phosphorylation by Aurora B modulates kinetochore disassembly in late anaphase. **a**. HeLa cells expressing Dsn1-WT-GFP were stained for DNA (blue), GFP (green), Nuf2 (red) and CENP-C (gray). Where indicated, cells were treated with 10 µM ZM447439 for 15 min prior to fixation. **b**. HeLa cells expressing Dsn1-EE-GFP were treated and stained as in (a). Scale bar, 5μm. **c**. In anaphase cells treated as in (a) and (b), Nuf2 at kinetochores was quantified as a function of distance from the central spindle. Using linear regression, for 17 Dsn1-WT-GFP cells, slope = −2.9 and is non-zero (p < 0.0001; F test). For 17 ZM447439-treated Dsn1-WT-GFP cells, 20 Dsn1-EE-GFP cells, and 16 ZM447439-treated Dsn1-EE-GFP cells, the slopes are −0.3, 0.03 and −0.1, respectively, which are not significantly different from zero (p = 0.29, 0.86 and 0.60). Comparing Nuf2 in Dsn1-WT-GFP cells with and without ZM447439, and comparing Nuf2 in Dsn1-WT-GFP with Dsn1-EE-GFP cells, the slopes are significantly different from one another (p < 0.0001, F test). Confidence intervals (95%) shown as fine lines. **d**. In telophase cells treated as in (a) and (b), Nuf2 was quantified at kinetochores. Coloured symbols show the results for individual kinetochores, black circles show the means for individual cells, and red bars show the means of these means. For Dsn1-WT-GFP cells +/-ZM447439, n = 9 and 12; and for Dsn1-EE-GFP cells +/-ZM447439, n = 11 and 7. ** adjusted p = 0.0054; **** adjusted p < 0.0001; by one-way ANOVA followed by Dunnett’s multiple comparisons test.

### Aurora B maintains kinetochore function in anaphase

To monitor kinetochore-microtubule attachments in anaphase, we turned to Astrin, a marker of stable end-on kinetochore-microtubule attachments^37^. The localization of Astrin to kinetochores is modulated by KMN components^38-40^. Astrin becomes localized at kinetochores as microtubules attach in prometaphase, and it clearly decorates kinetochores in metaphase and into anaphase^41,42^. Imaging of HeLa cells expressing Astrin-GFP^42^ and CENP-B-Cherry showed that Astrin-GFP remained detectable at kinetochores during anaphase until the chromosomes are in close proximity to the spindle poles, when Astrin appeared to be fully released from kinetochores (Figure 6a, Movie 5). In cells treated with Aurora B inhibitor, chromosomes moved more slowly away from the midzone^27,43^ and, in several cases, did not approach as closely to the spindle poles. In these cells, Astrin was lost from kinetochores closer to the midzone than in control cells, coinciding with the termination of chromosome movement prior to reaching the spindle poles (Figure 6b, Movie 6). Furthermore, in untreated cells with spontaneously lagging or bridging chromosomes, Astrin-GFP appeared to persist on kinetochores closer to the midzone (Figure S5a, Movie 7).

**Figure 6:**
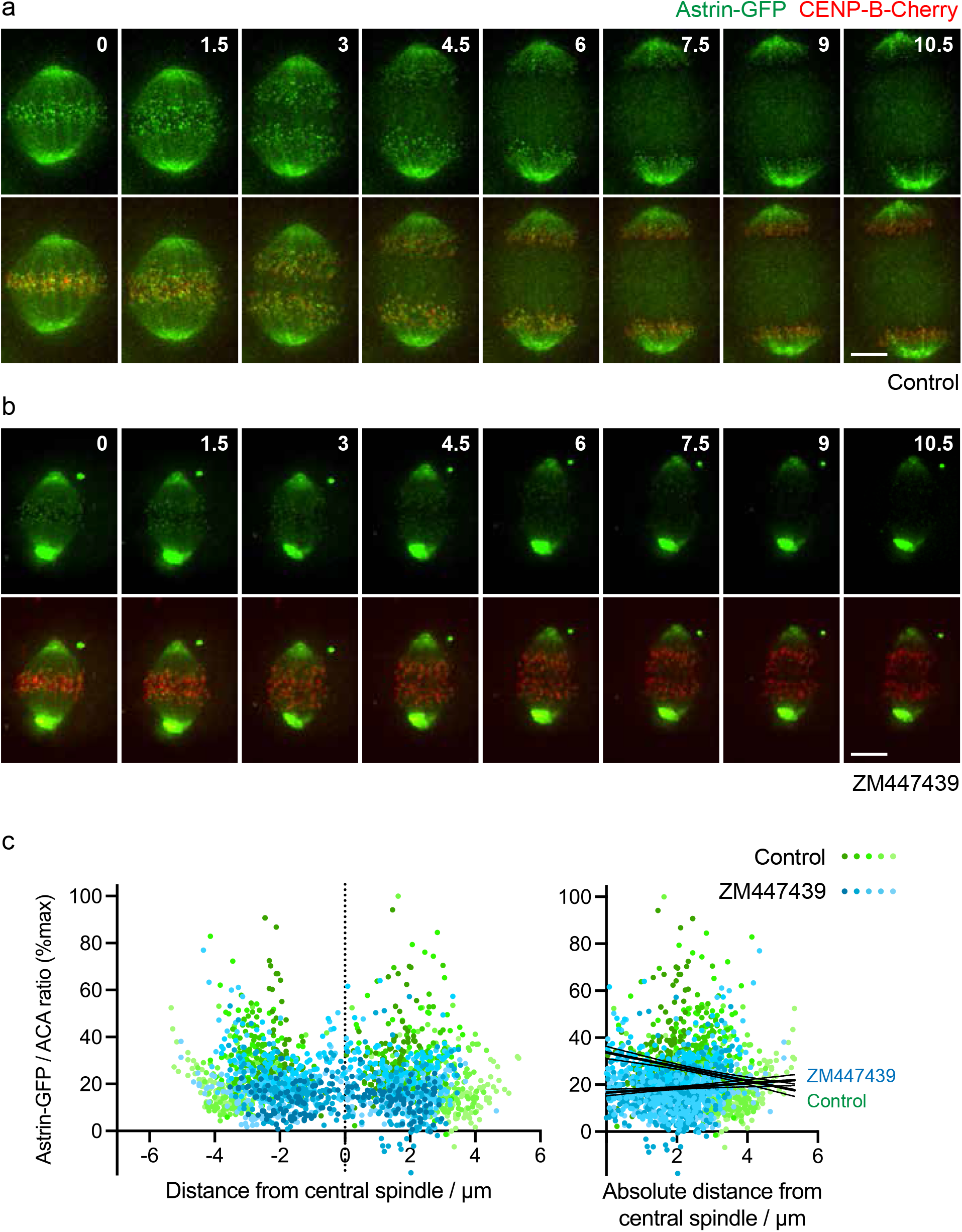
Astrin localization is influenced by Aurora B in anaphase. **a**. A HeLa cell stably expressing Astrin-GFP (green) and CENP-B-Cherry (red) was imaged every 30 s by iSIM super-resolution microscopy. **b**. A cell as in (a) was treated with 5 µM ZM447439 immediately prior to imaging, which began at time = 0. Scale bar, 5 µm. **c**. HeLa cells expressing Astrin-GFP were treated with 5 µM ZM447439 or control for 10 min, then cold treated to depolymerise labile microtubules, fixed and stained for DNA, GFP, and ACA. Astrin-GFP at kinetochores was quantified as a function of distance from the central spindle. Using linear regression, for 13 control cells, slope = −3.0 which is non-zero (p<0.0001, F test), and for 12 ZM447439-treated cells, slope = 0.99 which is non-zero (p = 0.0027; F test). The slopes are significantly different from one another (p < 0.0001, F test). Confidence intervals (95%) shown as fine lines.

In these live imaging experiments, the presence of Astrin on microtubules near the spindle poles^37^ introduced ambiguity into the timing of Astrin release from kinetochores. As an alternative method, we turned to immunofluorescence of cold-treated anaphase cells. Cold treatment favours disassembly of non-kinetochore microtubules in anaphase^44^, and we noticed that it reduced the intensity of Astrin-GFP on the spindle, allowing clearer visualization of kinetochore Astrin. Cold treatment may also reveal the influence of Aurora B on the stability of kinetochore attachments by favouring the disassembly of unstable kinetochore microtubules. Using this approach, we confirmed that Astrin-GFP levels on anaphase kinetochores were decreased by Aurora B inhibitor treatment (Figure 6c). Similar results were obtained in MPS1-IN-1-treated cells (Figure S5b,c). Clearly Aurora B influences the function of other spindle components^45^, but these results are consistent with the idea that Aurora B activity is required to maintain the stability and function of kinetochores as they move away from the midzone in anaphase.

## Discussion

The use of artificial centromere-targeted substrates previously revealed a gradient of Aurora B activity centered on the spindle midzone in anaphase^25^. Here we show that phosphorylation of a natural kinetochore substrate (Dsn1 S109) is sensitive to its distance from midzone Aurora B, suggesting that the Aurora B gradient modulates the activity of kinetochores themselves during anaphase. Prior to anaphase, phosphorylation of Dsn1 S100 and S109 by centromeric Aurora B promotes the association of Mis12C with its centromere receptors, CENP-C and CENP-T^3,5-7,12,19-21,46^. Here we show that Aurora B also regulates Dsn1 localization at kinetochores in anaphase. Specifically, midzone Aurora B-mediated phosphorylation of S100/S109 reduces the rate at which Dsn1 is lost from kinetochores as anaphase progresses. Loss of Aurora B activity also leads to premature loss of Astrin, a marker of stable microtubule attachments, from kinetochores. Therefore, the anaphase gradient of Aurora B serves to prolong the stability of kinetochores at a time when the loss of centromeric Aurora B might otherwise lead to premature kinetochore disassembly.

It is remarkable that structural components of the kinetochore such as Dsn1 appear to be shed as chromosomes are moving to the poles in anaphase, but this phenomenon has been reported for a number of KMN proteins^9,47^, and microtubule attachments appear relatively insensitive to the amount of Mis12C or functional Ndc80/Hec1 at kinetochores^48,49^. Nevertheless, it is possible that shedding represents continued remodeling of the kinetochore to provide dynamics optimal for anaphase chromosome movements. Indeed, it is notable that chromosomes slow down as anaphase progresses^13^, perhaps reflecting changes in kinetochore function and microtubule dynamics. In addition, we find that the timely loss of Aurora B-mediated phosphorylation on Dsn1 S100/S109 is required for the normal dissociation of KMN proteins from kinetochores at the end of anaphase.

It is also striking that the spatial gradient of total Dsn1 levels at anaphase kinetochores is shallower than that of Dsn1 S109ph, and that Dsn1 appears to remain at kinetochores for a time even when Dsn1 S109 is no longer phosphorylated, and when any possible Cdk1-dependent localization of Mis12C is expected to be low (see below). This may be due to a previously unexplained feature of Dsn1 regulation: that S100/S109 phosphorylation does not directly enhance binding to CENP-C (and CENP-T), but rather it displaces an autoinhibitory loop of Dsn1 to reveal the CENP-C binding region^7,12,19-21^. This mechanism means that the initial stable binding of Mis12C to kinetochores is highly dependent on phosphorylation of Dsn1 S100/S109 and requires high local Aurora B activity^46^, but that dephosphorylation does not necessarily lead to the immediate release of Dsn1. This would allow Mis12C to persist at kinetochores as Aurora B activity declines to maintain kinetochore function (Figure 7), a model supported by the reported high affinity of Mis12C binding to CENP-C^20^. We speculate that this property of autoinhibitory mechanisms may be important in this and other cellular systems.

**Figure 7:**
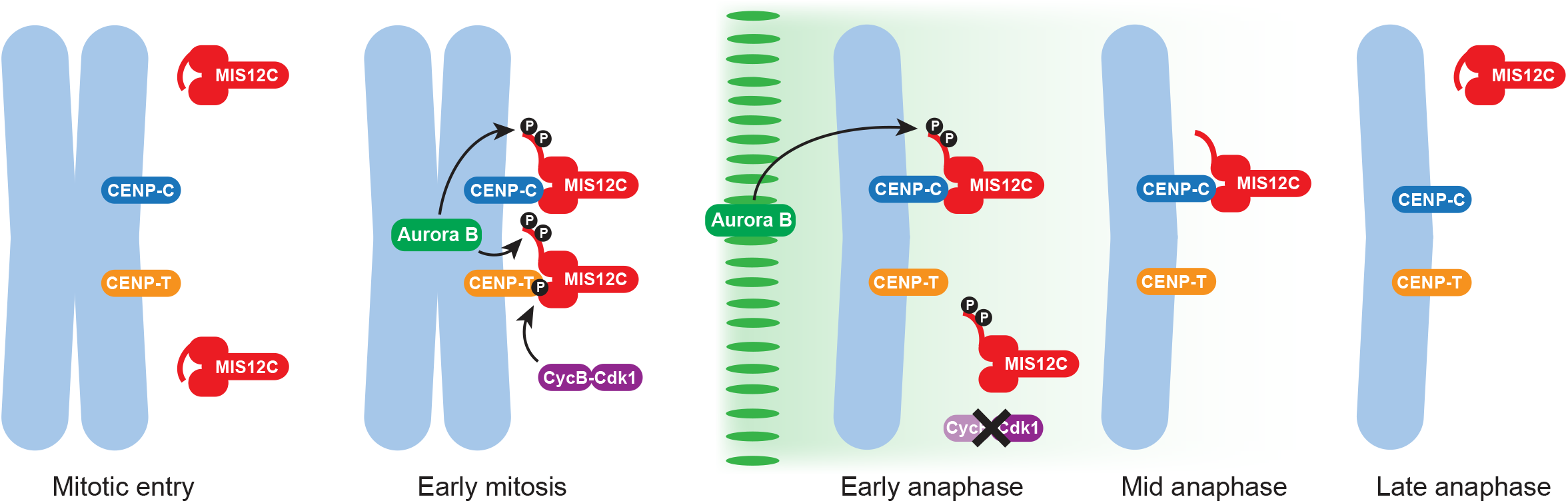
Model for Mis12C regulation at anaphase kinetochores. In early mitosis, phosphorylation at S100/S109 by Aurora B (green) displaces the basic region of Dsn1 and allows Mis12C (red) binding to CENP-C (blue) and CENP-T (orange). Phosphorylation of CENP-T by Cyclin B-Cdk1 (purple) is also required for Mis12C to bind CENP-T. In early anaphase, Cyclin B degradation and loss of Cdk1-dependent CENP-T phosphorylation releases Mis12C from CENP-T, while Aurora B gradient activity can prolong phosphorylation of Dsn1 S100/S109, allowing Mis12C retention on CENP-C. As chromosomes move away from the central spindle, declining Aurora B activity allows dephosphorylation of Dsn1 S100/S109, although full release of Mis12C is delayed because phosphorylation is not directly required for CENP-C binding. By late anaphase, Dsn1 returns to its autoinhibited conformation, and all Mis12C is released.

In early mitosis, Aurora B contributes to both the assembly of kinetochores (eg through Dsn1 S100/S109 phosphorylation), and to the destabilization of kinetochore-microtubule interactions to facilitate error correction (eg through phosphorylation of Ndc80/Hec1). This raises the question of whether the Aurora B gradient is able to destabilize microtubule attachments in anaphase, such as on the midzone-proximal regions of kinetochores of merotelic lagging chromosomes. While this model has some appeal, and we cannot entirely rule it out due to technical issues, we have been unable to demonstrate phosphorylation of Ndc80/Hec1 on Aurora target sites such as S44 and S55 in early anaphase (data not shown). Aurora B-mediated phosphorylation of Ndc80/Hec1 declines more strongly in metaphase than that of Dsn1^4,27-30^, suggesting that Dsn1 phosphorylation requires lower Aurora B activity levels than that of Ndc80/Hec1. Indeed, Dsn1 S100/S109 appear to be particularly good substrates for Aurora B *in vitro*, particularly once autoinhibition has been relieved^46,50^. It appears likely that the level of Aurora B gradient activity experienced by kinetochores in early anaphase is sufficient to maintain Dsn1 phosphorylation to stabilize kinetochores, but insufficient to destabilize kinetochore-microtubule interactions. Other substrates that are sensitive to the Aurora B gradient in anaphase include Histone H3S10^25,43^ and CENP-AS7 (this study), and we note that phosphorylation of Dsn1 may not be the only mechanism by which Aurora B influences kinetochore function in anaphase.

In previous work, Wurzenberger et al. found that the depletion of phosphatase regulators such as Sds22 and Repo-Man caused increased phosphorylation of Dsn1 S100 and transient pauses in anaphase chromosome migration^27^. At that time, it was not known that Dsn1 S100ph is involved in stabilizing kinetochores rather than in destabilizing microtubule attachments. In light of our study, it is perhaps more likely that failure to dephosphorylate Dsn1 compromises necessary changes in the structural properties of the kinetochore itself and so indirectly alters the dynamics required for chromosome movement. Alternatively, phosphatase depletion might allow other Aurora B targets such as Ndc80/Hec1 to be over-phosphorylated to destabilize attachments. Whether the phosphorylation of Aurora B substrates can reach levels high enough to destabilise microtubule attachments on lagging chromosomes in otherwise normal cells remains an open question.

As well as Aurora B, Cyclin B-Cdk1 plays a vital role in building kinetochores^8-12,21^, and our study is consistent with loss of Cdk1 activity also playing an important role in regulating kinetochore structure in anaphase (Figure 7). Indeed, before anaphase, up to 50% of Dsn1 molecules require phosphorylation of CENP-T by Cdk1 to be recruited to kinetochores^11,33^. It is clear that Dsn1 with phospho-mimetic residues at S100 and S109 is removed from kinetochores in early anaphase with kinetics similar to wild type Dsn1 (Figure 4c). Our two-phase decay model for loss of Dsn1 from kinetochores suggests that the Cdk1-mediated CENP-T-dependent population of Dsn1 dissociates with a half-life of approximately 5 min, consistent with the rapid dephosphorylation of many Cdk1-dependent sites in anaphase^8,35^. Non-degradable Cyclin B causes aberrant anaphase chromosome movements^22,51^, consistent with the need for timely dephosphorylation of substrates such as CENP-T. On the other hand, short-lived residual Cdk1 activity in early anaphase^34^ may sustain Dsn1 at kinetochores for a short time, influenced also by the rate of CENP-T dephosphorylation^9^. Together, Cdk1 and Aurora B co-operate to regulate Dsn1 localization and kinetochore structure in anaphase (Figure 7).

In summary, we have shown that phosphorylation driven by the Aurora B gradient helps sustain kinetochore structure over the time and distance necessary for normal anaphase chromosome segregation, and regulates kinetochore disassembly as cells enter telophase. This spatial regulation of kinetochore phosphorylation may also allow kinetochore stability on lagging chromosomes to be maintained to facilitate their movement to the poles in anaphase^52^.

## Acknowledgments

We thank Iain Cheeseman, Ulrike Gruneberg, Michael Lampson, Soonjoung Kim, and Hongtao Yu for gifts of antibodies, plasmids, and cell lines; Helder Maiato and Bernado Orr for discussions of unpublished work; Alex Laude, Rolando Berlinguer Palmini, Glyn Nelson and David Bulmer of the Newcastle Bioimaging Unit and Steven Coleman of Visitech for their invaluable help with microscopy; and Fangwei Wang for his contribution to the early stages of this work. This project was supported by BBSRC project grant BB/P020771/1, a Royal Society Wolfson Research Merit Award, and a Wellcome Investigator (106951/Z/15/Z) award to JMGH. For the purpose of open access, the author has applied a CC BY public copyright licence to any Author Accepted Manuscript version arising from this submission.

## Author Contributions

DP and ML designed and conducted experiments and analysed data. JMGH designed experiments, analysed data, and supervised the project.

## Methods

### Cells

HeLa Kyoto cells were grown in Dulbecco’s Modified Eagle’s Medium (DMEM; Sigma) with 5% (v/v) foetal bovine serum (FBS) and 100 U/ml penicillin-streptomycin. HeLa-TetOn cells expressing Dsn1-EGFP and Dsn1-EE-EGFP (a gift from Soonjoung Kim and Hongtao Yu, University of Texas Southwestern)^7^ were grown in similar conditions, with the addition of 150 µg/ml hygromycin B (Invitrogen). hTERT-RPE1 cells were grown in DMEM/F12 (Sigma) with 5% (v/v) FBS, and 100 U/ml penicillin-streptomycin. HeLa cells expressing Astrin-EGFP (a gift from Ulrike Gruneberg, University of Oxford)^42^ were grown in DMEM with 5% (v/v) FBS and 100 U/ml penicillin-streptomycin with 100 µg/ml G418. To produce a HeLa cell line stably expressing Astrin-EGFP and CENP-B-Cherry, this cell line was co-transfected with a 9:1 ratio of pCENP-B-Cherry (a gift from Michael Lampson; Addgene plasmid #45219) and pPGKpuro resistance plasmid. After selection at 2 µg/ml puromycin, two rounds of cloning by FACS was used to obtain a cell population with consistent moderate levels of Astrin-EGFP and CENP-B-Cherry expression. All cells were maintained in a humidified incubator at 37°C and 5% CO_2_.

### Cell treatments

For anaphase enrichment, in some experiments, cells were treated for 20 h with 2.5 mM thymidine (Sigma), washed 3 times in pre-warmed PBS (Sigma), twice in pre-warmed DMEM, and released for 10 h in 5% FBS/DMEM prior to fixation. To induce lagging chromosomes, HeLa and RPE1 cells were treated with 10 µM MPS1-IN-1 (MedChem Express) for 3 h prior to fixation. For Aurora B inhibition, cells were treated with 1 to 10 µM ZM447439 (Tocris) for 3 to 15 min. To depolymerise anaphase spindles, cells were first blocked in 1 µM nocodazole (Sigma) for 5 h and then, after release for 2 h, treated again with 10 µM nocodazole for 5 min prior to fixation. Dsn1-EGFP expression in HeLa-TetOn cells was induced using 1 µg/ml doxycycline (Sigma) for 20 h.

### RNA interference

For both Mklp2 and PRC1 depletion, HeLa cells suspended at 0.8 × 10^5^ cells/ml were transfected with 50 nM siRNA using HiPerfect transfection reagent (Qiagen) according to the manufacturer’s protocol and analysed after 48 h. The siRNAs used were: for Mklp2, 5’-CCACCUAUGUAAUCUCAUGTT-3’ (Integrated DNA Technologies); for PRC1, ON-TARGETplus siRNA SMARTpool L-019491-00-0005 (Dharmacon); and AllStars negative control siRNA 1027281 (Qiagen).

### Antibodies

For immunofluorescence analysis, rabbit polyclonal antibodies to Aurora B (Abcam ab2254, 1:1000), Nuf2 (Abcam ab122962, 1:500), Dsn1 (19.2B, 1:1000), Dsn1S109ph (19.2A or, where stated, 20.2A, 1:1000; Dsn1 antibodies were a gift from Iain Cheeseman, Whitehead Institute)^4^ and CENP-AS7ph (Upstate 07-232, 1:500); mouse monoclonal antibodies to Aurora B (AIM-1; BD Bioscience 611082, 1:100) and H3S10ph (6G3; Cell Signaling #9706, 1:500); guinea pig polyclonal antibodies to CENP-C (MBL PD030, 1:1000); chicken polyclonal antibodies to GFP (Applied Biological Materials G160, 1:1000); and human anti-ACA antibodies (Immunovision HCT-0100, 1:1000) were used. Secondary antibodies were donkey anti-rabbit and anti-mouse IgG Alexa Fluor Plus 594 (ThermoFisher A-32754 and A-32744), donkey anti-rabbit and anti-mouse IgG Fluor Plus Alexa 488 (ThermoFisher A-32790 and A-32766), goat anti-guinea pig and anti-human IgG Alexa 647 (ThermoFisher A-21450 and A-21445), goat anti-chicken IgY Alexa Fluor 488 (Thermofisher A-21449).

### Indirect Immunofluorescence

HeLa cells were grown on poly-L-lysine-coated coverslips (Corning BioCoat 354085). For Dsn1 (19.2B) and Dsn1 S109ph (19.2A/20.2A) staining, cells were pre-extracted in 120 mM PIPES, 50 mM HEPES, 20 mM EGTA, and 8 mM MgSO_4_ pH 7.0 (2x PHEM buffer) containing 1% Triton X100 for 5 min and then fixed in 4% (v/v) paraformaldehyde (ThermoFisher) in 2x PHEM buffer at 37°C. For Nuf2 staining, cells were fixed in ice-cold methanol (Fisher) for 10 min and then washed in PBS. For Astrin-EGFP staining, cold treatment was carried out by placing cells in L-15 medium (Sigma) with 20 mM HEPES on ice for 10 min, before fixation in 4% (v/v) paraformaldehyde in 1x PHEM buffer with 0.2% Triton X100 for 10 min at room temperature. Coverslips were incubated with 1% (v/v) BSA (Rockland) in PBS for 1 h at room temperature, and stained with primary antibody diluted in 1% BSA in PBS for 1 h at 37°C, and secondary antibodies for 45 min at 37°C. Coverslips were mounted on slides using ProLong Diamond with DAPI (P36962) or ProLong Glass with NucBlue (P36981) from ThermoFisher.

### Fluorescence microscopy

Images of fixed samples were acquired on a Leica SP8 confocal microscope equipped with a 63x 1.4 NA PlanApo Oil objective, capturing data at optimal Nyquist sampling using Leica LasX v3 software (Leica Microsystems, Germany), or on a Nikon A1 confocal microscope equipped with a 60x 1.4 NA PlanApo Oil objective, at optimal Nyquist sampling using Elements 5.22 software (Nikon, Japan), or on a Visitech VT-iSIM super-resolution Nikon TiE-based microscope (Visitech, UK) with instant SIM scanhead coupled to two Hamamatsu Flash4 v2 cameras (Hamamatsu, Japan) via a dual port splitter, using a 100x 1.49NA PlanApo Oil objective and Elements 5.21.03 software (Nikon, Japan).

Widefield images were captured with a Zeiss AxioImager microscope equipped with a 63x 1.4 NA PlanApo Oil objective using a Colibri1 LED light source, an AxioCam MrM camera, and ZEN 2.3 software (Zeiss, Germany). Optical sectioning was improved by using an Apotome 2 for structured illumination, capturing 3 images per focal plane and channel (Zeiss).

Live imaging was performed in glass-bottomed FluoroDishes (WPI) in Fluorobrite DMEM medium (ThermoFisher). DNA was stained with 25 nM SiR-DNA (Spirochrome), and ZM447439 was added at double the required final concentration in a volume equal to that of medium in the dish to ensure rapid mixing. Recording was started immediately after drug addition. Images were acquired on a Nikon A1R confocal microscope equipped with a 60x 1.4NA PlanApo Oil objective, at optimal Nyquist sampling using Elements 5.22 software (Nikon, Japan), or on the Visitech VT-iSIM super-resolution microscope as described above. A humidified environment at 37°C and 5% CO_2_ was maintained in an Okolab whole microscope and a stage-top incubators (Okolab, Italy).

### Image analysis

Fixed cell image quantification was performed using sum intensity projections in Fiji 2.1.0^53^. The midzone position was defined as the point equidistant between the edges of the chromosomes nearest the poles. Circular regions of interest (ROIs) were defined on the ACA or CENP-C channel, and the intensities within each ROI recorded for this and all other channels of interest. The mean of the intensities within 4 background ROIs placed in areas devoid of ACA or CENP-C, but within the chromatin mass, was then subtracted from the relevant channels. Staining intensity at each kinetochore is expressed as a ratio to the ACA or CENP-C intensity at that kinetochore. Dsn1-GFP levels at kinetochores in live cell imaging experiments were quantified using Imaris (Bitplane, Oxford Instruments), where the DNA channel was used to define the chromatin region within each cell for tracking in time. The intensity of the GFP signal within the defined volume was then recorded for each time point. The mean cytosolic background signal was subtracted from all data points. Normalisation, least squares linear regression, statistical analyses and data visualisation were carried out in Prism 9.0.1 (Graphpad).

For display purposes only, live images were deconvolved in Nikon Elements (Nikon, Japan) using the Richardson-Lucy algorithm. Where noted, fixed cell images were deconvolved with Huygens (Scientific Volume Imaging).

### Mathematical model

We assumed that there are two populations of Dsn1 associated with kinetochores, one associated with CENP-T and dependent on Cyclin B-Cdk1 (and Aurora B) that is removed rapidly in anaphase, and another associated with CENP-C, and dependent only on Aurora B, that is removed less rapidly in anaphase. The two-phase decay equation was then:

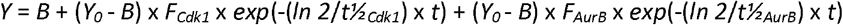

The parameters were: *Y* = amount of kinetochore-associated Dsn1; *B* = bottom plateau; *t* = time after anaphase onset; *Y*_*0*_ = Value of *Y* at *t* = 0, *F*_*Cdk1*_ = fraction of Dsn1 loss dependent on Cdk1; *F*_*AurB*_ = fraction of Dsn1 loss dependent on Aurora B; *t½*_*Cdk1*_ = half-life of Cdk1-dependent Dsn1; *t½*_*AurB*_ = half-life of Aurora B-dependent Dsn1. To fit the experimental data, we used least squares non-linear regression in Prism 9.0.1 (Graphpad), constraining *Y*_*0*_ = 1, *t½*_*Cdk1*_ to be equal in all conditions, and *t½*_*Cdk1*_ < *t½*_*AurB*_, and assuming that for Dsn1-WT, *F*_*Cdk1*_ = 0.5, *F*_*AurB*_ = 0.5, and for Dsn1-EE that *F*_*Cdk1*_ = 1, *F*_*AurB*_ = 0.

## Supplementary Figure Legends

**Figure S1: Dsn1S109ph and CENP-AS7ph show anaphase gradients in HeLa and RPE1 cells**.

**a**. HeLa cells treated with MPS1-IN-1 were stained for DNA (blue), Dsn1S109ph (green, antibody 20.2A), H3S10ph (red), and ACA (gray).

**b**. RPE1 cells treated with MPS1-IN-1 were stained for DNA (blue), α-Tubulin (red), CENP-AS7ph (green), and ACA (gray).

**c**. RPE1 cells treated with MPS1-IN-1 were stained for DNA (blue), Dsn1S109ph (green), and ACA (red). Channel brightness was adjusted individually for these two cells. Scale bars, 5μm.

**d**. RPE1 cells treated with MPS1-IN-1 were stained as in (c) and Dsn1S109ph was quantified at kinetochores of 5 cells as a function of distance from the central spindle. Using linear regression, slope = −2.7 and is non-zero (p < 0.0001; F test).

**e**. As for (d) but for 5 RPE1 cells not treated with MPS1-IN-1. Using linear regression, slope = −4.5 and is non-zero (p < 0.0001; F test). Confidence intervals (95%) shown as fine lines.

**Figure S2: The anaphase Dsn1S109ph gradient is compromised by short term Aurora B inhibition, or depletion of Mklp2 or PRC1**.

**a**. HeLa cells were treated as in Figure 2a, and stained for DNA (blue) and Aurora B (green) confirming that Aurora B is lost from the midzone upon nocodazole treatment.

**b**. HeLa cells were treated as in Figure 2b, and stained for DNA (blue) and Aurora B (green) confirming that (inactive) Aurora B remains at the central spindle upon short term ZM447439 treatment. Scale bars, 5 μm.

**c**. HeLa cells were treated with MPS1-IN-1 and then with 1 μM ZM447439 for 3 min prior to fixation and staining for DNA (blue), Dsn1S109ph (green), and CENP-C (red). Dsn1S109ph was quantified at kinetochores in 6 control and 5 ZM447439-treated cells. Using linear regression, for control cells, slope = −2.2 and is non-zero (p < 0.0001; F test). For ZM447439-treated cells, slope = 0.9 and non-zero (p = 0.0001). The slopes are significantly different from one another (p < 0.0001, F test).

**d**. As in (c), but for HeLa cells not treated with MPS1-IN-1. Dsn1S109ph was quantified at kinetochores in 7 control and 6 ZM447439-treated cells. Using linear regression, for control cells, slope = −3.5 and is non-zero (p < 0.0001; F test). For ZM447439-treated cells, slope = 0.004 and not significantly different from zero (p = 0.99). The slopes are significantly different from one another (p < 0.0001, F test).

**e**. HeLa cells were treated as in Figure 2d, but in the absence of MPS1-IN-1. Dsn1S109ph was quantified at kinetochores in 10 control and 8 Mklp2-depleted cells. Using linear regression, for control cells, slope = −2.9 and is non-zero (p < 0.0001; F test). For Mklp2-depleted cells, slope = −0.9 and is non-zero (p < 0.0001; F test). The slopes are significantly different from one another (p < 0.0001, F test). Confidence intervals (95%) shown as fine lines.

**f**. RPE1 cells were transfected with control or Mklp2 siRNA, treated with MPS1-IN-1, and stained for DNA (blue), Dsn1S109ph (green), Aurora B (red) and ACA (gray). Arrows indicate kinetochores with elevated Dsn1S109ph in control cells.

**g**. Following PRC1 depletion, the central spindle is disrupted, and Aurora B is retained at the equatorial cortex during anaphase. HeLa cells were transfected with PRC1 siRNA, fixed, and stained for DNA (blue), Dsn1S109ph or CENP-AS7ph (green), Aurora B (red) and CENP-C (gray).

Phosphorylation was retained only at those kinetochores in close proximity to Aurora B at the cell cortex (arrows). Deconvolved images are shown. Scale bars, 5 μm.

**Figure S3: Dsn1 is found in a gradient at anaphase kinetochores**.

**a**. HeLa cells enriched in anaphases by thymidine-release and treated with MPS1-IN-1 were stained for DNA (blue), total Dsn1 (green), and CENP-C (red). Scale bar, 5 µm.

**b**. Quantification of total Dsn1 (18 cells) at kinetochores as a function of distance from the central spindle. Using linear regression, slope = −2.0 and is non-zero (p < 0.0001; F test).

**c**. HeLa cells were treated with Mps1-IN-1 and then with 5 μM ZM447439 for 15 minutes prior to fixation and staining for DNA (blue), Dsn1 (green), and CENP-C (red). Dsn1 was quantified at kinetochores in 6 control and 5 ZM447439-treated cells. Using linear regression for control cells, slope = −2.6, and for ZM447439-treated cells, slope = −2.4, and both are non-zero (p < 0.0001; F test). The elevations or intercepts, but not slopes, are significantly different from one another (p < 0.0001, F test).

**d**. HeLa cells were treated with 5 μM ZM447439 for 5 minutes prior to fixation and staining for DNA (blue), Dsn1 (green), and CENP-C (red). Dsn1 was quantified at kinetochores in 6 control and 5 ZM447439-treated cells. Using linear regression, for control cells slope = −3.5, for ZM447439-treated cells slope = −3.8, and both are non-zero (p < 0.0001; F test). The elevations or intercepts, but not slopes, are significantly different from one another (p < 0.0001, F test). Confidence intervals (95%) shown as fine lines.

**e**. HeLa cells were untreated or treated with 5 μM ZM447439 for 5 minutes prior to fixation and staining for DNA (blue), Dsn1 or Dsn1S109ph (green), CENP-C (red) and H3S10ph (gray). Scale bars, 5 µm.

**Figure S4: Mathematical model of Dsn1 dissociation from anaphase kinetochores**.

**a**. The dissociation of Dsn1 from anaphase kinetochores was modelled as a two-phase exponential decay, assuming a starting condition of two molecules of CENP-T-associated Dsn1 and two molecules of CENP-C-associated Dsn1 per centromeric nucleosome ^36^, and a half-life for Cdk1-dependent Dsn1 dissociation of 5 min. The graph shows the effect of varying the half-life of Aurora B-dependent Dsn1 from 5 to 500 min.

**b**. Least squares linear regression fit of the two-phase exponential decay model to the data in Figure 4c (see Methods).

**Figure S5: Astrin localization in anaphase is influenced by Aurora B in MPS1-IN-1-treated cells**.

**a**. A living HeLa cell stably expressing Astrin-GFP (green) and CENP-B-Cherry (red) was imaged by iSIM super-resolution microscopy. Lagging kinetochores that maintain Astrin-GFP are indicated with white arrows.

**b**. MPS1-IN-1-treated HeLa cells expressing Astrin-GFP were exposed to 5 µM ZM447439 or control for 10 min, then cold treated to depolymerise labile microtubules, fixed and stained for DNA (blue), GFP (green), and ACA (red). Deconvolved images are shown. Scale bars, 5 µm.

**c**. For cells treated as in (a), Astrin-GFP at kinetochores was quantified as a function of distance from the central spindle. Using linear regression, for 19 control cells, slope = −2.2 which is non-zero (p<0.0001, F test), and for 11 ZM447439-treated cells, slope = 0.24 which is not significantly different from zero (p = 0.57; F test). The slopes are significantly different from one another (p < 0.0001, F test). Confidence intervals (95%) shown as fine lines.

**Movie 1**

Dsn1-WT-GFP (left panel) and DNA labeled with SiR-DNA (right panel) was imaged in a living HeLa cell at 1 min intervals. Deconvolved images are shown.

**Movie 2**

Dsn1-WT-GFP (left panel) and DNA labeled with SiR-DNA (right panel) was imaged in a living HeLa cell at 1 min intervals. 4 µM ZM447439 was added approximately 2 min prior to time = 0 min. Deconvolved images are shown.

**Movie 3**

Dsn1-EE-GFP (left panel) and DNA labeled with SiR-DNA (right panel) was imaged in a living HeLa cell at 1 min intervals. Deconvolved images are shown.

**Movie 4**

Dsn1-EE-GFP (left panel) and DNA labeled with SiR-DNA (right panel) was imaged in a living HeLa cell at 1 min intervals. 4 µM ZM447439 was added approximately 5 min prior to time = 0 min. Deconvolved images are shown.

**Movie 5**

A HeLa cell expressing Astrin-EGFP (green) and CENP-B-Cherry (red) was imaged every 30 s. Deconvolved images are shown. See also Figure 6a.

**Movie 6**

A HeLa cell expressing Astrin-EGFP (green) and CENP-B-Cherry (red) was imaged every 30 s. 5 µM ZM447439 was added approximately 2 min prior to time = 0 min. Deconvolved images are shown. See also Figure 6b.

**Movie 7**

A HeLa cell expressing Astrin-EGFP (green) and CENP-B-Cherry (red) with lagging kinetochores was imaged every 30 s. Deconvolved images are shown. See also Figure S5a.

