## Supplementary figures and images for "The Aurora B gradient sustains kinetochore stability in anaphase"

### Supplemental Figures

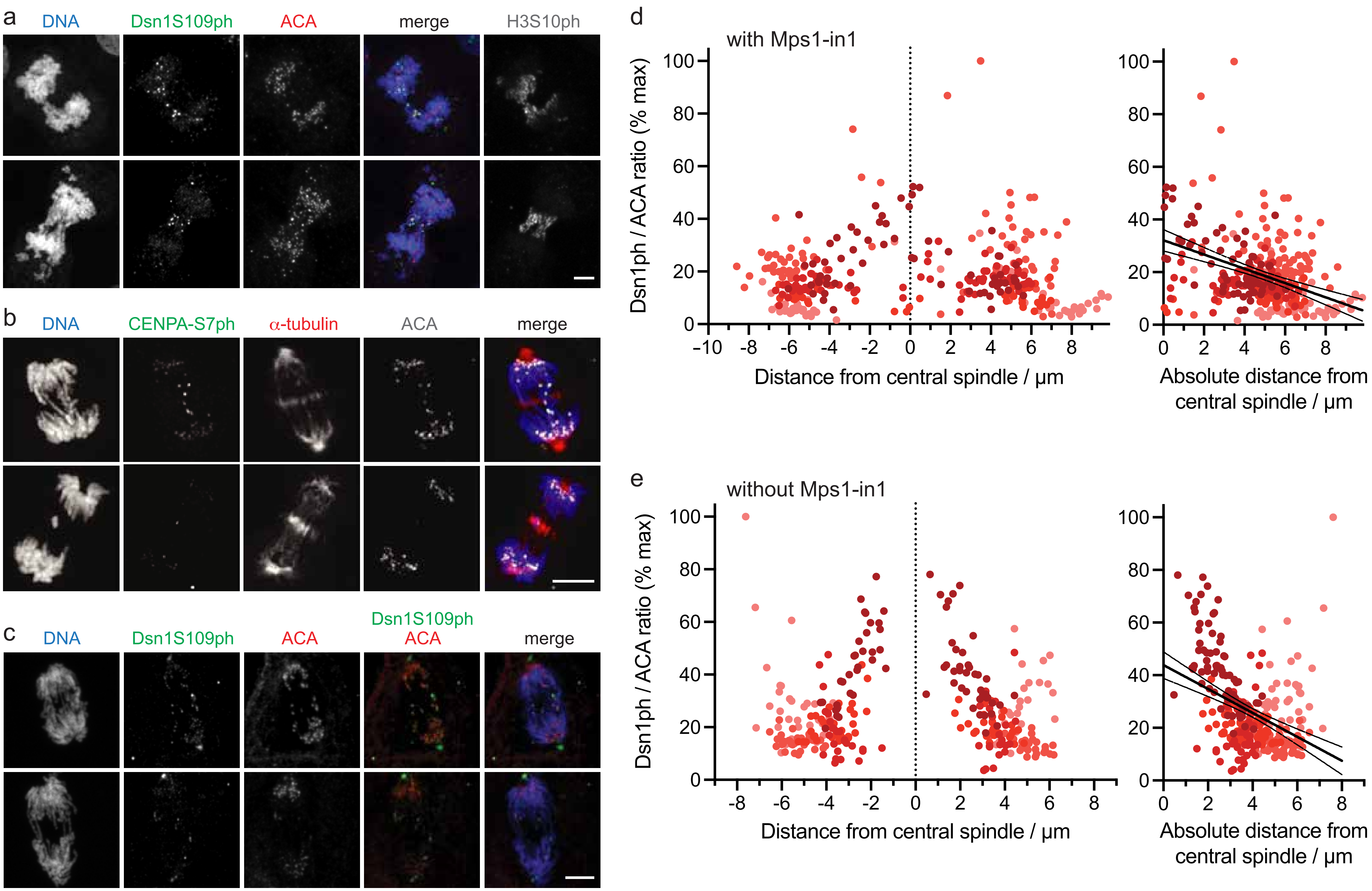

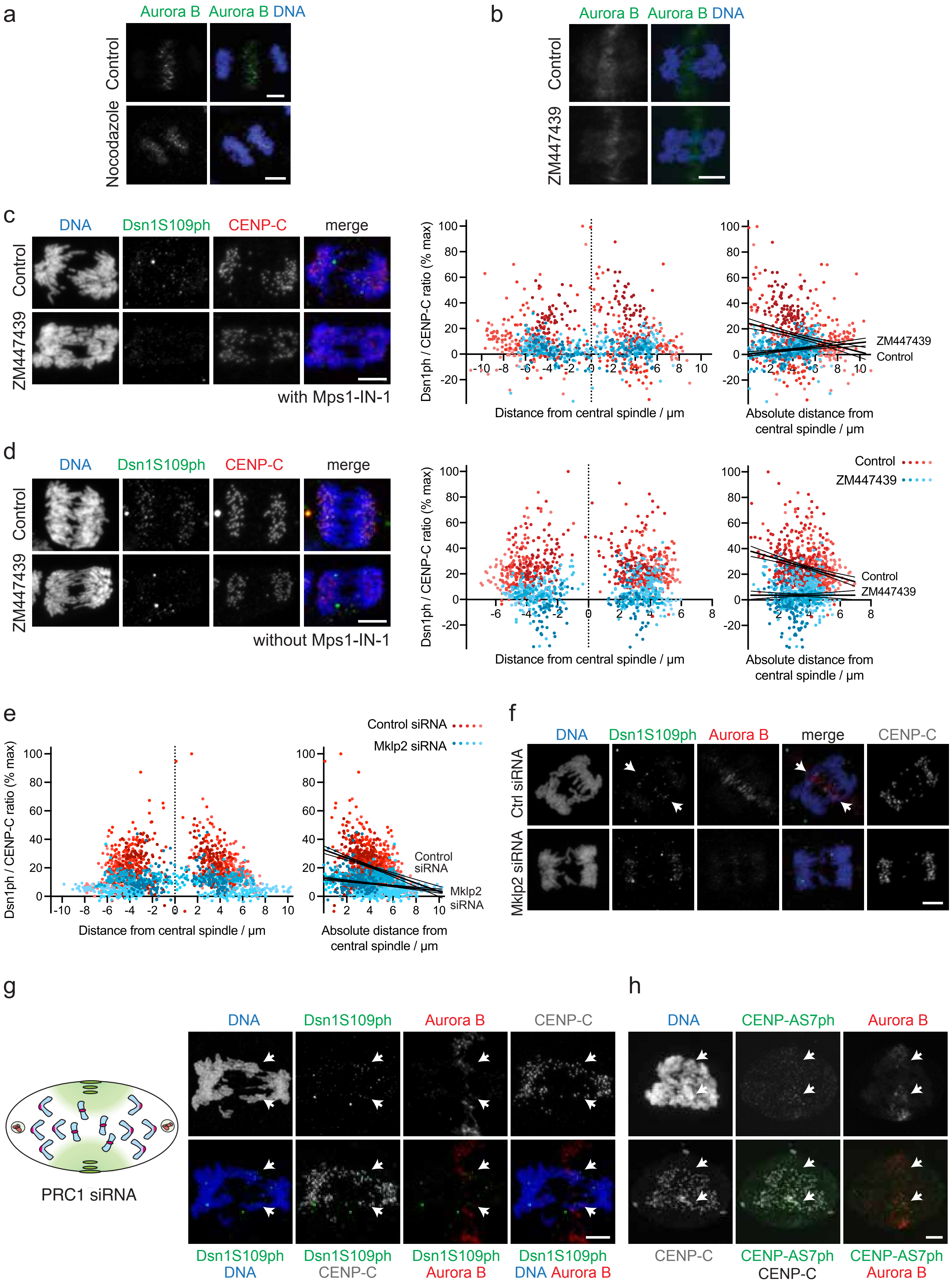

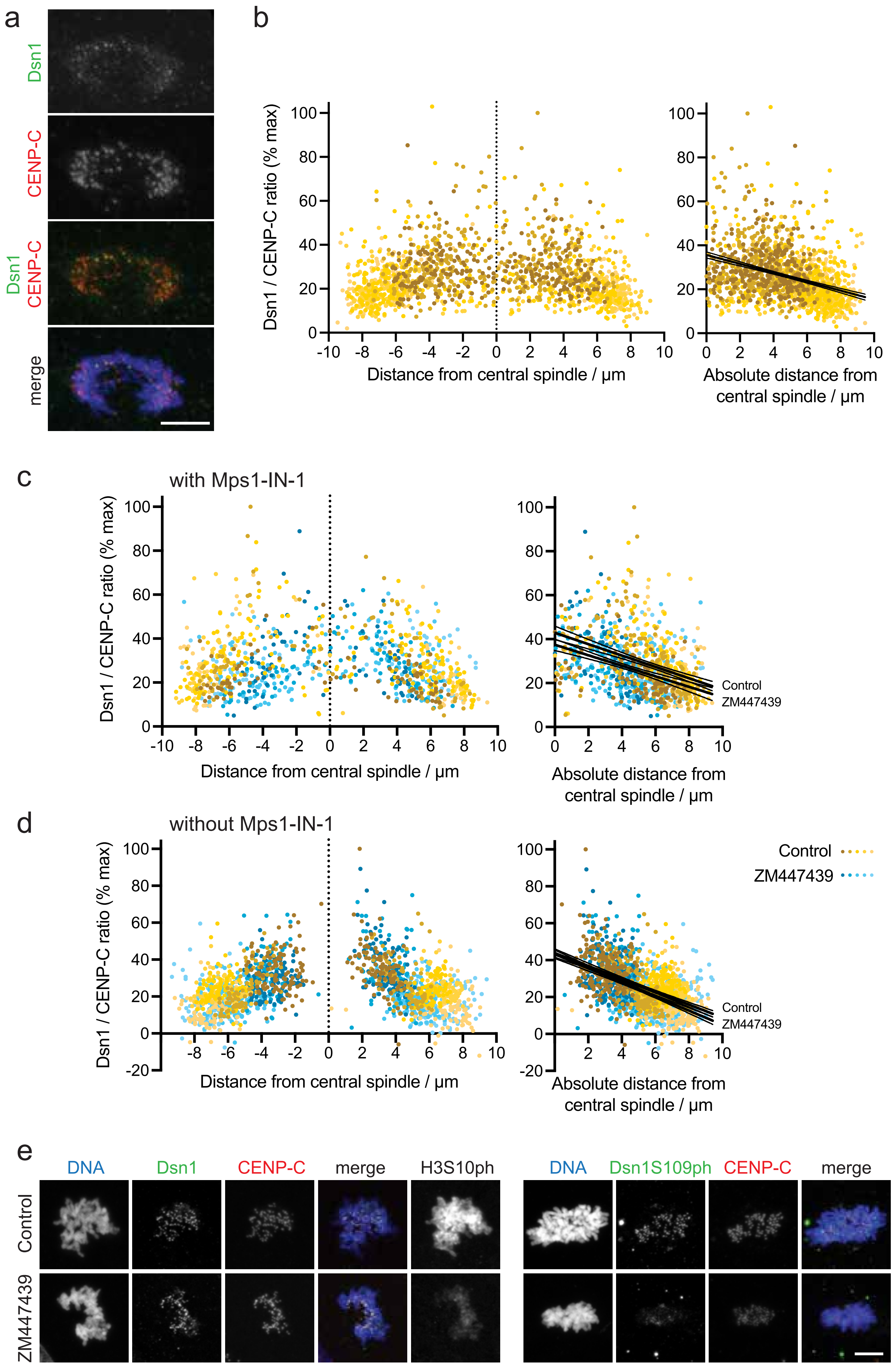

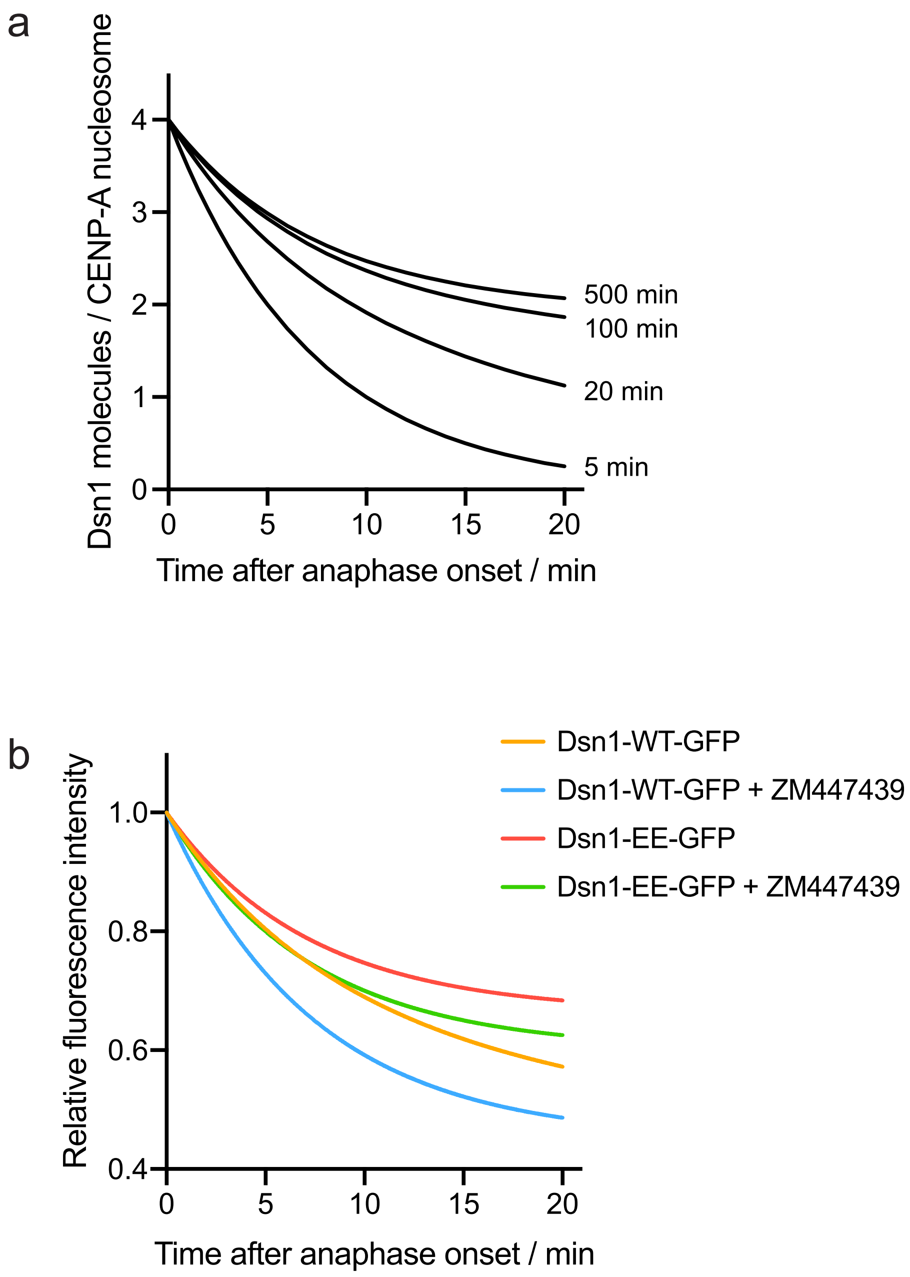

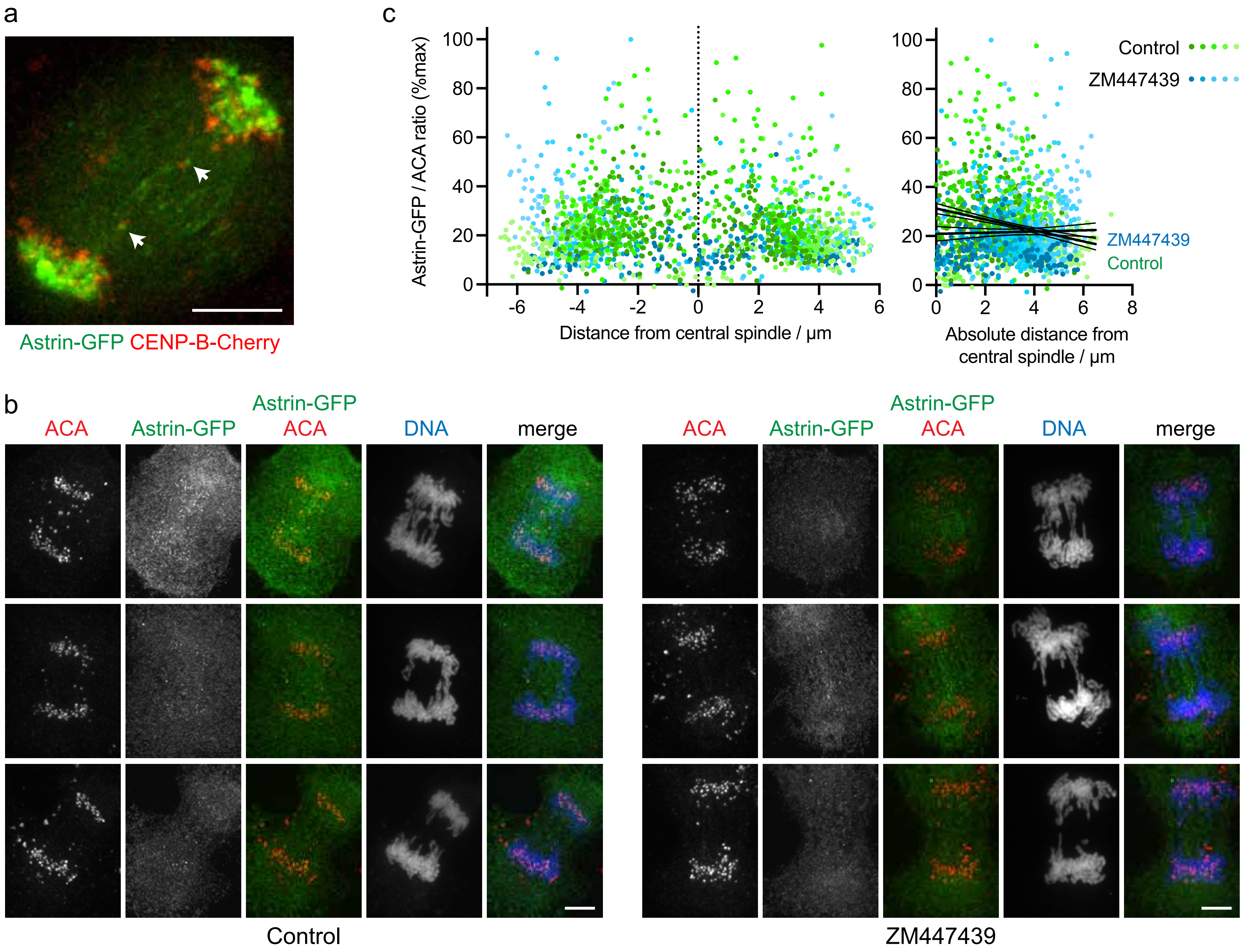
